# Which waters hydrate best? A study using brine-shrimp cysts (*Artemia franciscana*)

**DOI:** 10.1101/2020.09.23.310326

**Authors:** Tao Ye, Gerald H. Pollack

**Affiliations:** Department of Bioengineering, University of Washington, Box 355061, Seattle, WA, 98195, United States

**Keywords:** Brine shrimp, Water, Hydration

## Abstract

Hydration plays a particularly important role in health maintenance and general well-being. A wide assortment of drinking waters are currently available on the market. However, their ability to hydrate may vary. For studying hydration, a useful organism may be the cysts of brine shrimp. Those cysts may remain dehydrated and functionless for years, but regain function once hydrated. In this study, we first determined the optimal factors for assessing hydration in the brine-shrimp model, including aeration method and flow rate, salinity, and temperature. Various kinds of water, including tap water, bottled water, and water containing health-promoting agents, were tested by using this new method to evaluate their ability to hydrate. Tap water showed weak hydration, while some bottled waters (e.g., Kirkland Signature Purified Water) hydrated more effectively. Mineral water, such as Fiji water, was found to be a desirable option to maintain adequate and lasting hydration.

**Graphic abstract:** 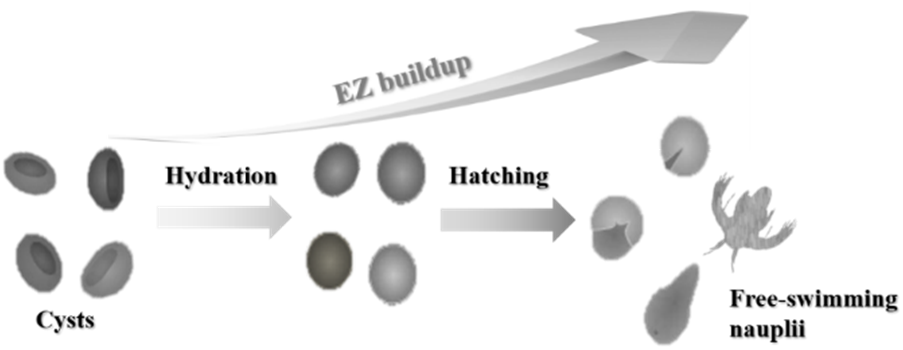

## 1. Introduction

Water is vital to all life. It is the main constituent of cells, tissues, and organs, and constitutes from 55 % body weight in the elderly, to 75 % in infants (Casado et al. 2015). It transports nutrients, oxygen, and waste products, regulates body temperature, and serves as a lubricant and shock absorber (Jéquier and Constant 2010, Tansey and Johnson 2015). Despite its well-established importance, many unresolved questions remain about this most essential component of our body and health (Popkin et al. 2010). For example, how does the type of water ingested affect human performance and health?

People often drink water as a result of water deficit, which is manifested as thirst (Jéquier and Constant 2010). Water intake maintains body hydration status. The role of water and hydration has been well-studied, particularly during physical activity (Murray 2007, Casa et al. 2000, Montain et al. 1999, Michael N. Sawka and Timothy D. Noakes 2007). Dehydration can significantly impair exercise performance, reducing endurance, increasing fatigue, altering thermoregulatory capability, reducing motivation, and increasing perceived effort (Michael N. Sawka and Timothy D. Noakes 2007, Montain and Coyle 1992, Cheuvront et al. 2003). Rehydration can reverse these deficits, restoring physiologic function (Paik et al. 2009). The faster the rehydration, the better the sports performance, especially for athletes.

The physical characteristics of rehydration beverages, including salinity, color, sweetness, temperature, flavor, carbonation, and viscosity, appear to influence hydration (Casa et al. 2000). Previous literature has reported studies on several popular rehydration beverages, including caffeinated diet-cola, carbohydrate-electrolyte solution, coconut water, sodium-enriched coconut water, and sports drinks (containing glucose and sodium ion [Na^+^]), etc. (González-Alonso et al. 1992, Saat et al. 2002, Ismail I 2007, Maughan et al. 2015). Drinks containing the highest electrolyte (primarily sodium and potassium) contents seem to be most effective at maintaining hydration. However, a number of results have been reported, some contradictory (Ismail I 2007, Maughan et al. 2015), leaving the question of what kind of water is the best for hydration and maintaining hydration status.

While many kinds of waters are widely accessible, most previous studies have focused on sports drinks. Due to the complexity of assessing rehydration and performance, those studies have generally involved tedious processes, which have resulted in large errors (Maughan and Shirreffs 2010). Therefore, it has seemed necessary to develop a convenient and precise method to assess the effects of different waters (e.g., tap water and bottled water) on hydration.

Brine shrimp (genus *Artemia*), commonly known as sea monkeys, have the intriguing property that their dry embryos may be reversibly hydrated and dehydrated (Mamontov 2017, Morris 1971). These embryos are enclosed in protective shells, and are known as cysts. Appropriate hydration can initiate metabolic activity and macromolecular synthesis, whereas dehydration can reversibly terminate embryonic development (Mamontov 2017, Morris 1971). Brine shrimp have been considered useful organisms for studies on hydration/dehydration. Previous research has explored the kinetics of hydration/dehydration in those specimens and the impacts of successive hydration/dehydration cycles (Morris 1971, El-Magsodi et al. 2016, El-Magsodi et al. 2014, Wang et al. 2010). These reports encouraged us to employ brine shrimp to study the impacts of different types of water on hydration. There are multiple advantages associated with using brine shrimp in bioassay: cost-effectiveness; availability; and easy manipulation and maintenance under laboratory conditions. Those features make brine shrimp a promising method for assessing water hydration (Libralato et al. 2016).

This study aims to address the question: which water is best for hydration? In this study we determine the factors that contribute to brine shrimp hydration and develop an optimal method for assessing their hydration. Various kinds of water, including tap water, bottled water, and water containing health-promoting agents, were tested by using established and optimized methods to evaluate their capacity to hydrate.

## 2. Experimental Section

### Materials

Technical-grade brine-shrimp cysts, *Artemia franciscana*, of Great Salt Lake origin with a hatch rate of 80 % were purchased from Brine Shrimp Direct, Utah, USA. All cysts were in the quiescent dry state and stored in a moisture-free container at 4 °C in a refrigerator before use. Sodium chloride (NaCl) was obtained from Fisher Scientific. Probiotics (Culturelle Kids Packets Daily Probiotic Supplement) were bought from Culturelle. Tulsi extract (Holy Basil) was obtained from Herb Pharm (Williams, Oregon). Bottled waters of different brand were purchased from their merchandisers without any further treatment. Deionized water (DI water, 18.2 MΩ cm) was collected from a Barnstead D3750 Nanopure Diamond purification system.

### Water sample preparation

Commercially available sea salt blends were not used to avoid the presence of unknown compounds in the mixtures. Instead, we optimized a hydration method by using saline (NaCl) solutions prepared in DI water. A range of saline solutions (0 - 0.63 M) were tested. Artificial seawater was made by adding 37 g of sodium chloride in 1 L of DI water (0.63 M), which made its salinity equal to ∼ 36 Practical Salinity Unit (PSU). On average, seawater in the oceans has a salinity of approximately 35 PSU, which means there are 35 g of salts in 1 L of seawater.

Water containing probiotics and Tulsi extract were prepared in DI water as described in our previous publication.(Sharma et al. 2018) Tap water was collected from the laboratory faucet and allowed to sit for 12 h before experiments. Fountain water was collected from a fountain in the laboratory building and allowed to sit for 12 h before experiments. Boiled water in the experiments was prepared by boiling tap water or fountain water and then allowing it to cool to room temperature.

### Hydration procedures

Detailed hydration procedures are as follows:

1. Saline solution was preheated to a certain temperature (20, 25, or 35 °C) in a 1 L Erlenmeyer flask (PYREX™ VISTA™ narrow mouth, Corning 709801L) on a hotplate (Thermo Scientific™ HP88857100).
2. After said temperature was reached, 0.4 g of cysts were added to the saline solution. The mixture was then kept under continuous stirring or aeration for 48 h with the room light on. Aeration was supplied from the flask bottom to keep the cysts well mixed.
3. After 12 h of hydration, 3 sub-samples of 1 mL each were withdrawn every 4 h from the suspension and the number of live brine shrimp was counted. 3 sub-samples were withdrawn and counted for each time point to obtain a mean number of live brine shrimp and its standard deviation. 1 mL of the suspension was withdrawn using a 1 mL pipette and then divided into small drops on a Petri dish so that the live brine shrimp could be observed with the naked eye. Brine shrimp that could move or swim during observation were considered live or fully hydrated, while those that did not show any movement after 15– 30 s of observation were considered dead or unhatched. We define *cyst hydration number* (CHN) as the number of live brine shrimp measured in 1 mL of the suspension.
4. No food was supplied to the brine shrimp during the experiments. Each experiment was conducted twice. Error bars in the figures show standard deviations calculated from these repeated experiments.

## 3. Results and Discussion

Several factors have been reported to impact the hydration and hatching of brine shrimp cysts. They include salinity, aeration, temperature, pH, and illumination (Libralato et al. 2016). To establish the optimal method, we investigated variations of several of those factors, including aeration-flow rate and method, stirring speed, salinity, and temperature.

### Effect of aeration flow rate

Oxygen is an absolute requirement for cyst development. Aeration through a diffuser is generally believed to sustain large quantities of aquatic organisms (e.g., fish and shrimp) by producing small air bubbles and increasing dissolved oxygen (DO) (Boyd 1998). Thus, the effect of aeration with an air stone at different flow rates on the hydration of brine shrimp cysts was first studied.

As shown in **Figure 1**, fully hydrated brine shrimp that can move or swim did not emerge until around 16 h regardless of the aeration flow rate. At 16 h, cyst hydration numbers (CHNs) of 0.7 ± 0.6, 25.7 ± 2.1, and 15.7 ± 3.2 were measured for aeration flow rates of 0.1, 0.5, and 1 L min^−1^, respectively. The CHNs at 16 h for 0.5 and 1 L min^−1^ were significantly greater than that of 0.1 L min^−1^, especially for 0.5 L min^−1^. It is apparent from these results that a moderately high flow rate can substantially expedite hydration. The transition of cysts to the emerged nauplius that can swim free requires sufficient supply of energy (Clegg 1964). According to literature, the aerobic oxidation of trehalose is the major source of energy for this transition. Therefore, we expected that anything that adequately reduces trehalose oxidation will slow the emergence of free-swimming nauplius (i.e., the first free-swimming period of brine shrimp) (Clegg 1964). Flow rate of 0.1 L min^−1^ may deliver deficient DO for trehalose oxidation, resulting in insufficient energy for the cysts to develop (Clegg 1964, Sorgeloos and Persoone 1975).

**Figure 1.**
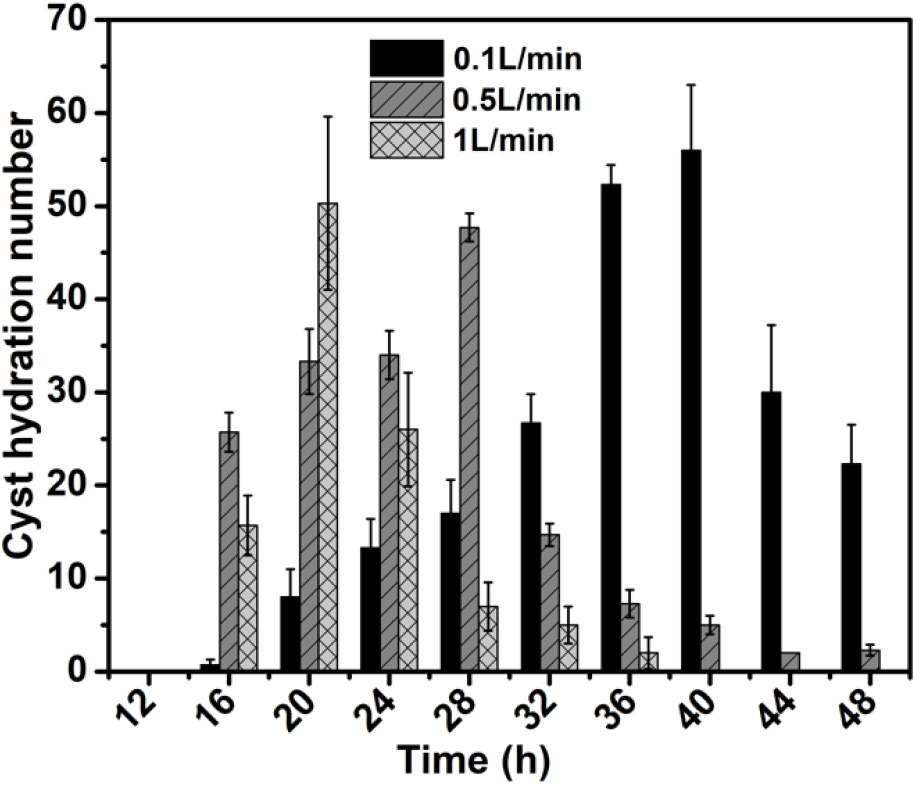
Effect of aeration flow rate on brine shrimp cyst hydration. The vertical axis shows the cyst hydration number (CHN), which is the number of live brine shrimp measured in 1 mL of suspension withdrawn from the hatching flask. The air was distributed to the saline solution containing brine shrimp cysts through an air stone diffuser (Model: ASC-016, Pawfly^®^) and provided from the bottom to keep the suspension well circulated. The air flow was controlled by a gas flowmeter (Part number: VIVIAN197, JIAWANSHUN). One end of the flowmeter connected to the air stone, the other end connected to the laboratory air supply nozzle outlet through plastic tubing. Experimental conditions: solution temperature, 25 °C; NaCl concentration, 0.63 M; cyst loading, 0.4 g in 1 L of solution.

For 0.1 L min^−1^, the CHN increased progressively from 16 h, reached its maximum value of 56.0 ± 7.0 at 40 h, and then gradually decreased. A similar trend was observed for 0.5 and 1 L min^−1^ except that their maximum CHNs of 47.7 ± 1.5 and 50.3 ± 9.3 were reached faster, at 28 and 20 h, respectively. This result further confirms that high aeration flow rates facilitate cyst hydration. However, these maximum CHNs are close to each other and their differences are statistically insignificant (*P* > *0*.*05*). This indicates that aeration flow rates merely impact the maximum cyst hydration. Other factors may limit optimal hydration.

In comparison to air stones, we did another experiment bubbling air through a 5 mL-pipette tip, which produced larger air bubbles than air stones. Results are shown in **Figure 2**. The CHNs showed a similar trend to that of air-stone bubbling in that they increased progressively from 16 h, reached their maximum values, and then gradually decreased. The decrease of CHNs after maximization was probably due to the lack of food in the suspension since no food was supplied.

**Figure 2.**
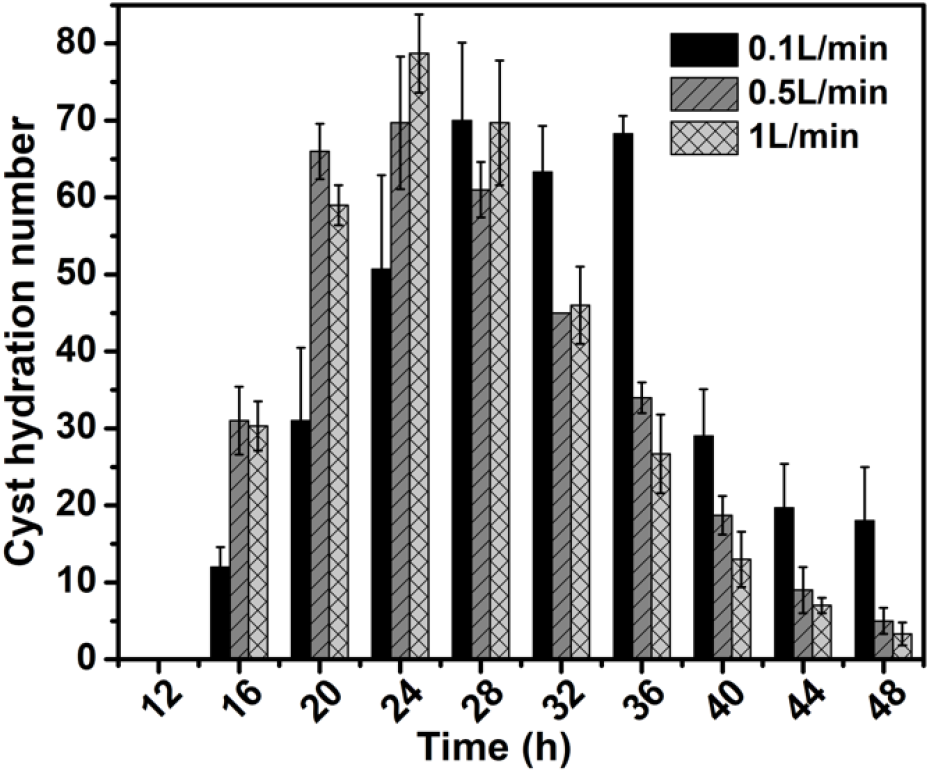
Effect of aeration flow rate on brine shrimp cyst hydration. The air was distributed to the saline solution containing brine shrimp cysts through a 5 mL-pipette tip (Fisherbrand™ Maxi Pipet Tips). The wider end of the pipette tip was connected to the gas flowmeter through plastic tubing, while the smaller end was used as a gas diffuser, bubbling air to the suspension. Experimental conditions: solution temperature, 25 °C; NaCl concentration, 0.63 M; cyst loading, 0.4 g in 1 L of solution.

The data shows that bubbling air through the 5 mL-pipette tip considerably improves cyst hydration. At 16 h, CHNs of 12.0 ± 2.6, 31 ± 4.4, and 30.3 ± 3.2 were measured for aeration flow rates of 0.1, 0.5, and 1 L min^−1^, respectively (**Figure 2**). Pipette tip bubbling also significantly enhanced maximum cyst hydration vs. air stone bubbling. The maximum CHNs of 68.3 ± 2.3, 69.7 ± 8.6, and 78.7 ± 5.1 were reached at approximately 36, 24, and 24 h for 0.1, 0.5, and 1 L min^−1^, respectively. These maximum CHNs are appreciably larger than those obtained with air stone bubbling (*P < 0*.*05*). Air stone bubbling seemed to retard cyst hydration. CHNs at each sampling point in pipette tip bubbling (**Figure 2**) were notably greater than those of air stone bubbling (**Figure 1**, except at 40 h for 0.1 L min^−1^).

The above results imply that pipette-tip bubbling is more favorable for cyst hydration than air stone bubbling. The thick cyst shell surrounding the embryo plays an important role in its osmotic properties (Wang et al. 2010). Air stone bubbling produces a fine mist of bubbles, which may damage the cyst shells and thus deteriorate their hydration and hatching. These small bubbles may also clog the shrimp’s feeding system and starve them. Therefore, pipette tip bubbling was used in our following experiments.

### Effect of stirring

Adequate aeration enables the cysts to hydrate and hatch. Aeration not only provides DO but also keeps the cysts well circulated in the suspension, both of which are beneficial to cyst hydration (Sorgeloos and Persoone 1975). The enhanced cyst hydration with high aeration thus could be ascribed to sufficient DO, circulation, or both. In order to ascertain how circulation contributes to cyst hydration, a series of experiments under different stirring conditions were conducted. We investigated different stirring speeds of 0, 80, and 400 rpm with magnetic stirring. To compare magnetic stirring with mechanical stirring, we also conducted an experiment using a home-made overhead stirrer, which minimized the impact of magnetic field that can be generated by magnetic stirring, and potentially improve cyst hydration (Yuriy et al. 2010, Sagdilek et al. 2013). Results are shown in **Figure 3**.

**Figure 3.**
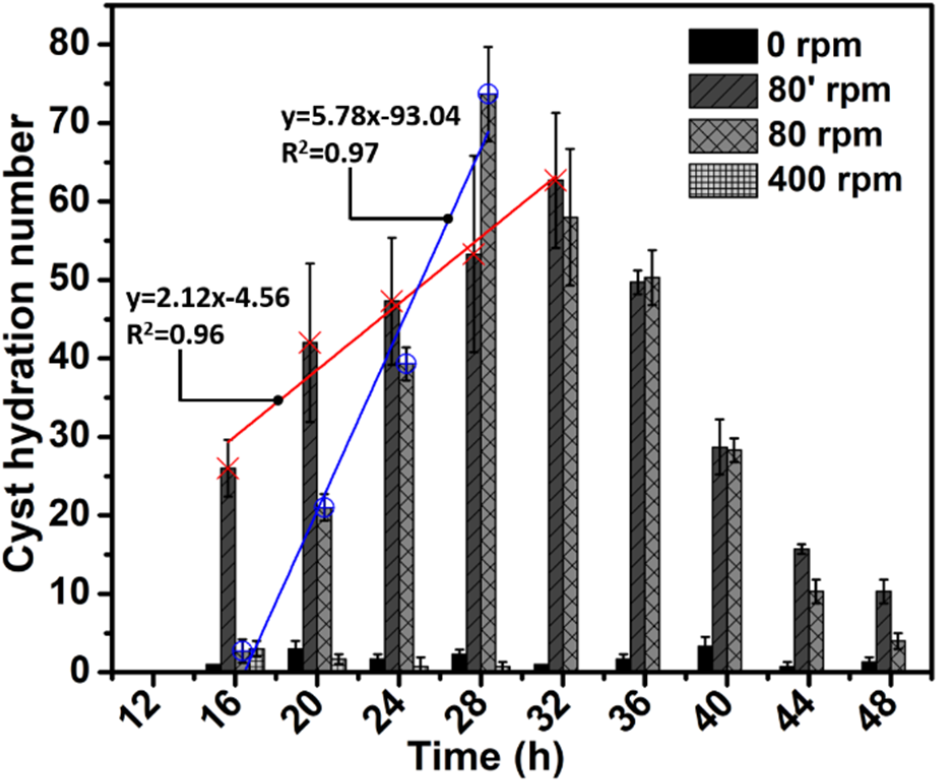
Effect of stirring on brine shrimp cyst hydration. Experiments at 0, 80, and 400 rpm were carried out on a magnetic hotplate. 80’ rpm indicates overhead stirring, which was performed on a hotplate with a home-made overhead stirrer (without magnetic stirring). Experimental conditions: solution temperature, 25 °C; NaCl concentration, 0.63 M; cyst loading, 0.4 g in 1 L of solution.

Mechanical stirring by the overhead stirrer appears to promote cyst hydration. At 16 h, the CHN of 80 rpm by overhead stirring reached 26.0 ± 3.6, which is considerably higher than those of 0, 80, and 400 rpm observed with magnetic stirring. The home-made overhead stirrer has a larger swept area (swept diameter, ∼16 mm) than the magnetic stir bar (Diameter, 8 mm; Length, 50.8 mm; Fisherbrand™), which leads to stronger water circulation. Water circulation can improve DO in the flask by mixing DO-supersaturated surface water with deeper water of lower DO concentration (Boyd 1998). Therefore the 80 rpm of overhead stirring likely enhanced DO in the suspension by providing stronger water circulation than with 80 rpm of magnetic stirring.

Interestingly, the CHN of 80 rpm by magnetic stirring increased rapidly after 16 h and attained its maximum of 73.7 ± 6.0 at 28 h, while the CHN of 80 rpm by overhead stirring gradually reached its maximum value of 62.7 ± 8.6 at 32 h. Though the difference in these maximum values was statistically insignificant (*P* > *0*.*05*), the CHN of 80 rpm by magnetic stirring after 16 h increased markedly faster (slope = 5.78, blue line, **Figure 3**) to its maximum than that of 80 rpm by overhead stirring (slope = 2.12, red line, **Figure 3**), indicating that magnetic stirring may be conducive to cyst hydration. Yuriy et al. suggested that magnetic field-induced chromatin granule formation in *Artemia* nauplia could promote dormant cyst activation (Yuriy et al. 2010). The effect of magnetic field on cyst hydration is beyond the scope of this study but worthy of further investigation.

It should also be noted that both 0 and 400 rpm largely limited cyst hydration. A few live brine shrimp were observed from 16 h to 48 h for 0 rpm. For 0 rpm, cysts accumulated at the bottom of the flask and were subjected to anaerobic conditions, which interrupted cyst development (Sorgeloos 1980, Clegg 2012). The CHNs of 400 rpm were also small and gradually decreased from 16 h to 28 h. After 28 h, no live specimens were observed. Stirring at 400 rpm may be too intensive and cause damage to cyst shells (Clegg 2005).

Overall, the role played by stirring may primarily be mixing DO-supersaturated surface water with deeper water of lower DO concentration (Boyd 1998). When comparing stirring (**Figure 3**) with aeration (**Figure 2**), aeration appears to facilitate cyst hydration. A significant number of live brine shrimp was observed even for a low aeration flow rate of 0.1 L min^−1^. Though low speed stirring (80 rpm, both magnetic and mechanical) could result in similar maximum CHNs to those of aeration, stirring generally reached these maximum values later (around 28–32 h) than aeration (at 24 h, for 0.5 and 1 min^−1^ L).

### Effects of salinity and temperature

According to the literature, the period between immersion of the cysts in the solution and the emergence of the embryos protruding from the shell is referred to as the ‘*lag period*’ (Clegg 1964). Afterwards, the hatching membrane ruptures and a free-swimming nauplius is born (Clegg 1966). Free-swimming brine shrimp were not observed until after 12 h for all the above experiments (**Figures 1–3**). It has been observed that the time of appearance of swimming brine shrimp occurred in a similar time range (∼12 h) as our observation in a previous study in which cysts were subjected to several hydration-dehydration cycles (Morris 1971). It is apparent that factors other than aeration and stirring determine the lag period.

Salinity and temperature have been reported to be the most significant abiotic factors that influence cyst development (Clegg 1964, Michael., von Hentig 1971, Jennings and Whitaker 1941, Arun et al. 2017, Browne and Wanigasekera 2000). Hence, we further investigated the effects of salinity and temperature on cyst hydration. Results are shown in **Figure 4**. Both salinity and temperature were found to be important factors affecting cyst hydration.

**Figure 4.**
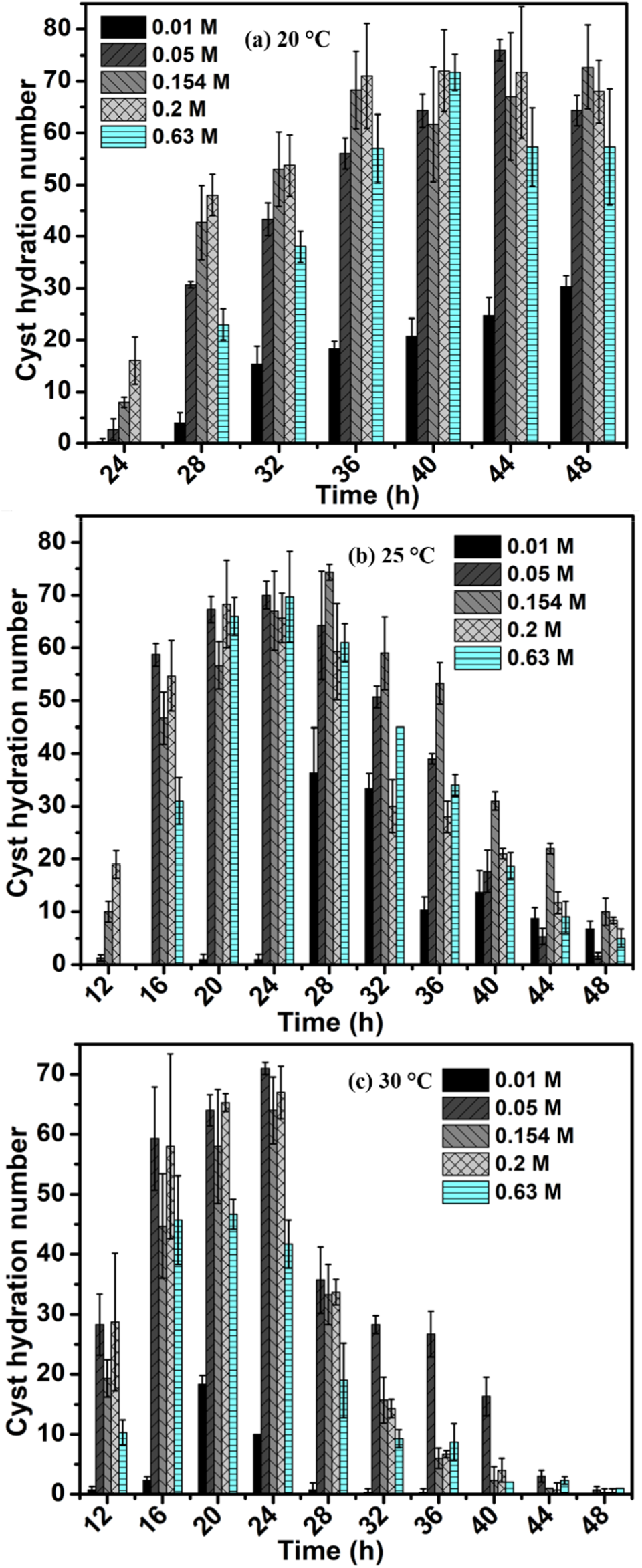
Effects of salinity and temperature on brine shrimp cyst hydration. The air was distributed to the saline solution containing brine shrimp cysts through a 5 mL-pipette tip. Experiments were conducted on a hotplate, which controlled the solution temperature. (a) 20 °C; (b) 25 °C; (c) 30 °C. Other experimental conditions were the same and kept as follows: aeration flow rate, 0.5 L min^−1^; cyst loading, 0.4 g in 1 L of solution. NaCl concentrations of 0.01, 0.05, 0.154, 0.2, and 0.63 M are equivalent to 0.55, 3, 9, 12, and 36 PSU, respectively.

Temperature higher than 30 °C and lower than 20 °C were not investigated because temperature above 32 °C is highly stressful for *Artemia* embryo development and temperature lower than 20 °C can significantly slow and delay cyst hatching (Arun et al. 2017). Except for 20 °C, the CHNs all showed an increasing and then decreasing trend with increasing time (**Figures 4b** and **c**), which was similar to what was observed in the above experiments (**Figures 1–3**). At 20 °C, the CHNs gradually increased from 24 h to 48 h (**Figure 4a**). Although we did not continue our experiments after 48 h, the CHNs were about to reach their maxima at around 48 h, and would probably decline after 48 h since no food was supplied in our experiments. We thus expect that the CHNs at 20 °C would also show a similar increasing and then decreasing trend if we further continued our experiments.

The increase in temperature from 20 to 30 °C substantially shortened the lag period from 24 h to 12 h. However, temperature change did not noticeably impact the maximum CHNs, which were all around 70 in these experiments. In terms of the experiments at 30 °C (**Figure 4c**), it seems that there was a minimum time of about 12 h for the lag period, before which time no free-swimming brine shrimp were observed. We did not observe any swimming brine shrimp at 8 h (data not shown). Similar results were obtained by Morris and coworkers who found that cysts could enter a latent period of about 12 – 18 h before the embryo emerged (Morris 1971). The lag period was found to be 10 h in the study by El-Magsodi et al., which is slightly shorter than our results.(El-Magsodi et al. 2014) However, it is highly possible that there may be some live brine shrimp at 10 h since a number of live brine shrimp were observed at 12 h in our experiments. Overall, it is apparent from these results that the major effect of temperature was on lag period rather than maximum CHNs.

Cyst development depends on the state and rate of hydration and can be manipulated by external salinity (Dana and Lenz 1986). The effect of different salinities on CHNs was not statistically significant among medium salinities (i.e., 0.05, 0.154, and 0.2 M), which is in agreement with the findings by Arun et al.(Arun et al. 2017) It has also been reported that optimal CHNs can be generally obtained in a salinity range of 0.08 – 0.60 M (Lavens 1996). However, too low salinity (i.e., 0.01 M) significantly reduced CHNs. The CHNs obtained in 0.01 M experiments were mostly smaller than those of other salinities. No live brine shrimp were observed in experiments with pure DI water without any addition of NaCl (data not shown).

High salinity results in a decrease in the effective concentration of water in the solution and thus can impact the hydration level of the cyst (Dana and Lenz 1986). As shown in **Figures 4a** and **b**, delays in hatching were observed for the highest salinity experiments (i.e., 0.63 M) as no free-swimming nauplius was found at the first sampling times (i.e., 24 h in **Figure 4a** and 12 h in **Figure 4b**).

### Evaluation of different water on cyst hydration

Based on above experiments, we established the optimal experimental conditions for cyst hydration and hatching (i.e., pipette-tip aeration with a flow rate of 0.5 L min^−1^; temperature 30 °C; salinity 0.2 M). We then tested various kinds of water, such as tap water and bottled water, by using these optimized conditions to evaluate their effects on hydration. Quality parameters for the water tested are listed in **Table 1**. Results are shown in **Figure 5**.

**Table 1.**
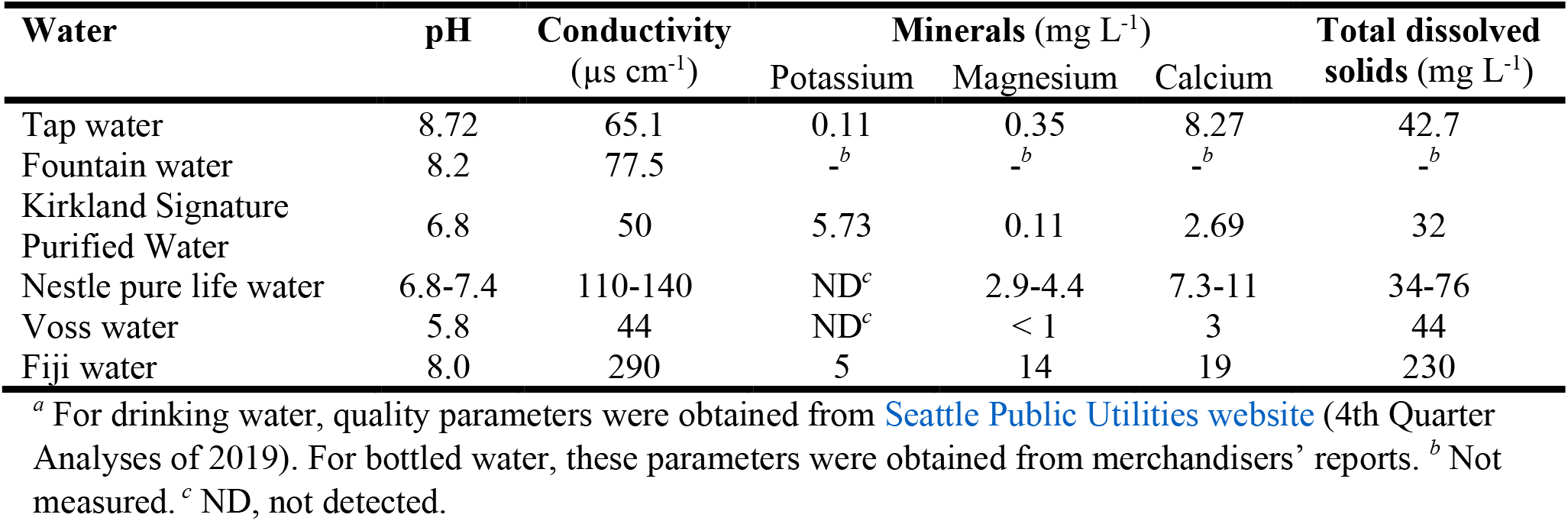
Quality parameters for the various water tested in our experiments.^*a*^

| Water | pH | Conductivity<br>($\mu\text{S cm}^{-1}$) | Minerals (mg L<sup>-1</sup>) | | | Total dissolved<br>solids (mg L<sup>-1</sup>) |
| --- | --- | --- | --- | --- | --- | --- |
|  |  |  | Potassium | Magnesium | Calcium |  |
| Tap water | 8.72 | 65.1 | 0.11 | 0.35 | 8.27 | 42.7 |
| Fountain water | 8.2 | 77.5 | <sup>-b</sup> | <sup>-b</sup> | <sup>-b</sup> | <sup>-b</sup> |
| Kirkland Signature<br>Purified Water | 6.8 | 50 | 5.73 | 0.11 | 2.69 | 32 |
| Nestle pure life water | 6.8-7.4 | 110-140 | ND<sup>c</sup> | 2.9-4.4 | 7.3-11 | 34-76 |
| Voss water | 5.8 | 44 | ND<sup>c</sup> | < 1 | 3 | 44 |
| Fiji water | 8.0 | 290 | 5 | 14 | 19 | 230 |

<sup>a</sup> For drinking water, quality parameters were obtained from Seattle Public Utilities website (4th Quarter Analyses of 2019). For bottled water, these parameters were obtained from merchandisers' reports. <sup>b</sup> Not measured. <sup>c</sup> ND, not detected.

**Figure 5.**
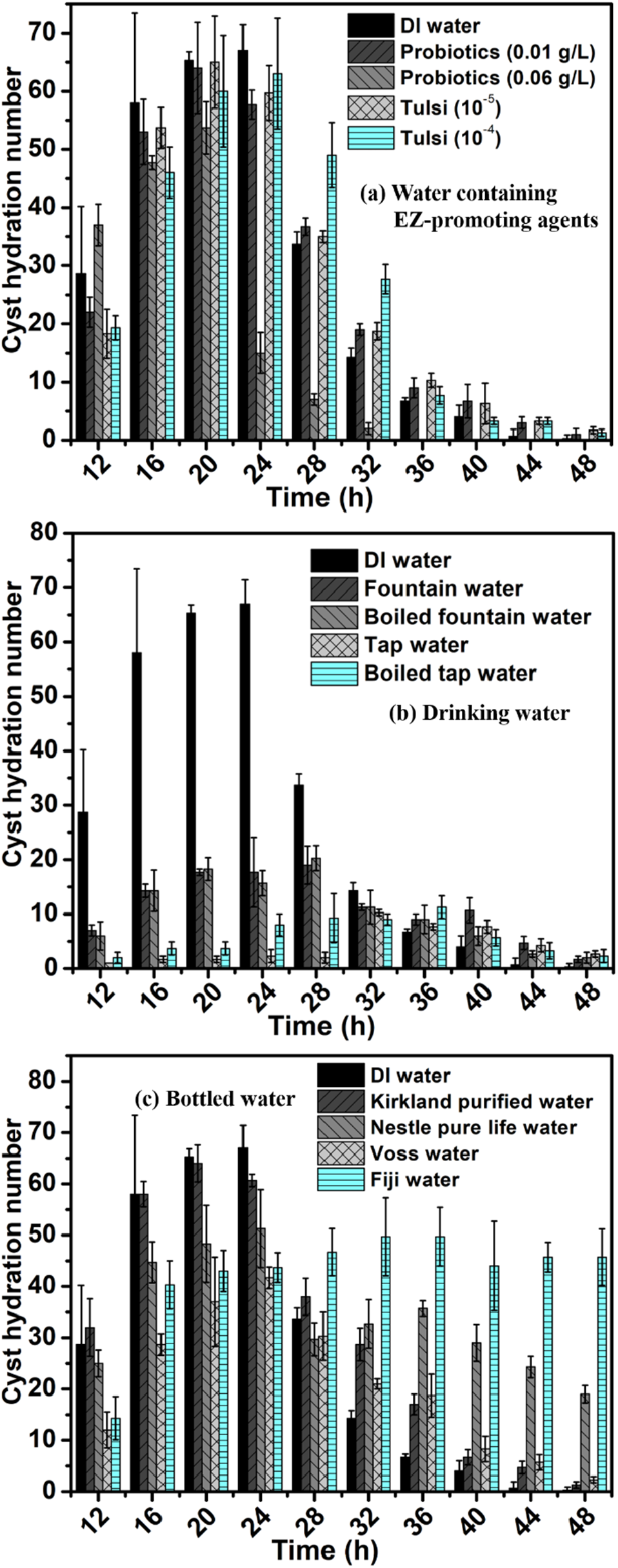
Effects of different waters on brine shrimp cyst hydration. Experimental conditions were as follows: aeration flow rate, 0.5 L min^−1^ (distributed through a 5 mL-pipette tip); temperature, 30 °C; cyst loading, 0.4 g in 1 L of solution. DI water was used as control. (a) Water containing EZ-promoting agents; (b) Drinking water; (c) Bottled water.

Cell water is mainly interfacial and ordered. This water has been termed “exclusion zone” (EZ) water (Sharma et al. 2018, Pollack 2013, Sharma and Pollack 2020). EZ water is critical to cellular health and function (Pollack 2001). Substances such as tulsi (holy basil) and probiotics known to enhance biological function have been shown to promote EZ water formation (Sharma et al. 2018). Therefore, we suspected that these substances could plausibly benefit an organism by building EZ water, thus hydrating cells, normalizing cell function, and restoring health (Sharma and Pollack 2020). By this mechanism, such substances would be deemed EZ promoting. We thus speculated that water containing these EZ-promoting agents might be able to enhance hydration and thus were the first to be tested in our experiments.

Comparatively, it appears that EZ-promoting agents did not improve CHNs (**Figure 5a**). Most CHNs obtained by water containing EZ-promoting agents were statistically similar to those of DI water. However, some observations deserve mention. High probiotic concentration (0.06 g/L) seemed to expedite cyst hydration and hatching since a high CHN value was observed at the first sampling point (i.e., 12 h). Though this CHN value was within the standard deviation compared to that of DI water, the CHN of 0.06 g/L probiotics reached its maximum value relatively fast at around 20 h. Curiously, after attaining its maximum value, it decreased markedly. Contrarily, the CHN of high tulsi concentration (10^−4^) increased gradually from 12 h, reached its maximum value around 24 h, and then declined relatively slowly. Overall, it is likely that a certain concentration of EZ-promoting agents in water would improve cyst hydration and hatching. We only tested two different concentrations of EZ-promoting agents to draw firm conclusions on this issue. Further study will be required to confirm or deny this speculation.

Since tap water and fountain water are major sources of public drinking water, we tested them. Considering that many people use boiled water, we also boiled these drinking waters and tested them. Interestingly, all of these drinking waters showed poor results compared to DI water (**Figure 5b**). *Before 32 h*, their CHNs were substantially lower than those of DI water. Meanwhile, CHNs of tap water was much lower than those of fountain water. Boiling tap water slightly enhanced CHNs, while boiling fountain water did not show any improvement. *After 32 h*, the difference among those CHNs were statistically insignificant.

Tap water has qualities similar to those of other waters, as shown in **Table 1**, except for its high pH. pH higher than 8 is actually beneficial for cyst hydration and hatching (Lavens 1996). Thus, these conventional water quality parameters are not able to account for the poor results obtained by tap water. Tap water usually contains disinfectants, such as chlorine and chloramines, which may prevent brine shrimp cysts from hatching. Seattle drinking water treatment utilities use chlorine. In our experiments, we let the tap water sit for 12 h and preheated it to 30 °C for another 12 h, which supposedly removed some free chlorine.

Chlorine disinfection leads to the formation of toxic disinfection byproducts, though at trace concentration levels (Sedlak and von Gunten 2011). Boiling tap water slightly improved the CHN results, which might be associated with partial removal of chlorine or these volatile byproducts (Zhang et al. 2015, Liu et al. 2020). Water fountain uses tap water as the source. The better results achieved by fountain water might have been due to the removal of chlorine and these volatile byproducts by the filters installed in the fountain (Saleh et al. 2001). Chlorine and toxic byproducts are health-impairing agents that can diminish the amount of EZ water (Casado et al. 2015, Sharma et al. 2018, Pollack 2013, Kundacina et al. 2016). In our previous work, we found several health-impairing agents (e.g., anesthetics and glyphosate) diminished EZ buildup (Sharma et al. 2018, Kundacina et al. 2016). We thus hypothesize that the poor results obtained with these types of drinking water (**Figure 5b**) could be due to the presence of these toxic compounds, curtailing EZ water formation.

We further tested some popular brands of bottled water (**Figure 5c**). Before 32 h, Kirkland Signature Purified Water (from Costco) and DI water both showed high CHNs and their results were comparable. After 32 h, the CHNs of Kirkland Signature Purified Water were mostly higher than those of DI water. It also deserves to be emphasized that the CHN of Kirkland Signature Purified Water reached its maximum faster (at 20 h) than that of DI water (at 24 h), indicating that Kirkland Signature Purified Water may have a better overall hydration result than DI water.

Similarly, Fiji water and Voss water show comparable results before 28 h, while the CHNs of Fiji water were significantly higher than those Voss water after 28 h. Interestingly, the results obtained with Nestle pure life water were somewhat between those obtained with Kirkland Signature Purified Water /DI water and Fiji water/Voss water. It is also worth noting that the CHNs of Fiji water gradually increased until 28 h and then reached a plateau, while the CHNs of all other water showed a decreasing trend after attaining their maximum. This profound effect of Fiji water in sustaining the further development of brine shrimp cysts after hydration and hatching is of significance and worthy of further investigation. Fiji water is artesian water containing a substantial amount of minerals and other ingredients that may offer health benefits, which we anticipate should promote EZ water formation (Maughan et al. 2015, Sharma et al. 2018, Pollack 2013).

Though DI water showed high CHNs before reaching its maximum at 24 h, its CHNs declined notably faster than those of other water. DI water theoretically has no minerals and organic materials, while other bottled waters all have certain amounts of minerals and dissolved solids (**Table 1**), which may be able to sustain the further development of cysts after hydration and hatching. This may also explain why Fiji water, which has an extraordinarily high content of minerals and dissolved solids, has such a profound effect on the further development of cysts after hydration and hatching. Therefore, mineral water, such as Fiji water, is an advisable option to maintain adequate hydration. Overall, DI water and Kirkland Signature Purified Water had a faster hydration effect since their CHNs were mostly higher than others, while Fiji water seems to have had a lasting hydration effect as its CHNs were maintained nearly constantly after reaching its maximum as early as 16 h.

## 4. Conclusions

Optimal experimental conditions for brine shrimp cyst hydration and hatching, including aeration method and flow rate (0.5 L/min with 5 mL-pipette tip bubbling), salinity (0.2 M), and temperature (30 °C), were established and used as a straightforward method for assessing hydration. Various kinds of water, including tap water, bottled water, and water containing EZ-promoting agents, were then tested by using this method to evaluate their effects on hydration. Tap water, which contained EZ-impairing agents (e.g., chlorine and disinfection byproducts), substantially reduced cyst hydration and hatching, while water containing EZ-promoting agents (e.g., probiotics and minerals) improved cyst development. These findings provide additional support to our hypothesis based on previously published observations that agents that benefit health build EZ.(Sharma et al. 2018, Sharma and Pollack 2020)

## Acknowledgements

We appreciate the funding from the Software AG (SAGST) Foundation, award number: CS-P12665, Germany. We also acknowledge useful scientific discussions with Abha Sharma, Alexis Traynor-Kaplan, Anqi Wang, Jianzhi Huang, Laura Colton, Rainier Stahlberg, Sally Landefeld, and Zheng Li. We did not receive any support from the bottled water brands that were used in the study and did not endorse them. The bottled water were randomly chosen from the market.

